# Hallucination proneness alters sensory feedback processing in self-voice production

**DOI:** 10.1101/2023.07.28.550971

**Authors:** Suvarnalata Xanthate Duggirala, Michael Schwartze, Lisa K. Goller, David E. J. Linden, Ana P. Pinheiro, Sonja A. Kotz

## Abstract

**Background:** Sensory suppression occurs when hearing one’s self-generated voice, as opposed to passively listening to one’s own voice. Quality changes of sensory feedback to the self-generated voice can increase attentional control. These changes affect the self-other voice distinction and might lead to hearing non-existent voices in the absence of an external source (i.e., auditory verbal hallucinations (AVH)). However, it is unclear how changes in sensory feedback processing and attention allocation interact and how this interaction might relate to hallucination proneness (HP).

**Study Design:** Participants varying in HP self-generated and passively listened to their voice that varied in emotional quality and certainty of recognition — 100% neutral, 60-40% neutral-angry, 50-50% neutral-angry, 40-60% neutral-angry, 100% angry, during EEG recordings.

**Study Results:** The N1 auditory evoked potential was more suppressed for the self-generated than externally generated voices. Increased HP was associated with (i) an increased N1 response to the self-compared to externally generated voices, (ii) a reduced N1 response for angry compared to neutral voices, and (iii) a reduced N2 response to unexpected voice quality in sensory feedback (60-40% neutral-angry) compared to neutral voices.

**Conclusions:** The current study highlights an association between increased HP and systematic changes of the emotional quality and certainty in sensory feedback processing (N1) and attentional control (N2) in self-voice production in a non-clinical population. Considering that voice hearers also display these changes, these findings support the continuum hypothesis. However, additional research is needed to validate this conclusion.

## Introduction

Sensations arise inevitably and incessantly from various internal and external sources. As we can predict the sensory consequences of self-generated actions, we suppress these sensations. For example, we perceive the sound of our own footsteps as less intense than those of another person. Accordingly, self- and externally-generated events differ in how we respond and adjust to them in a dynamic environment. The internal forward model provides a mechanistic explanation for such “sensory suppression”^1–3^. The model suggests that an internal copy of a motor plan (efference copy) is used to predict the sensory consequences of self-generated actions to prepare the brain for incoming sensory information. The perceived sensory feedback (reafference signal) is processed by comparison to this prediction, resulting either in a match or a mismatch (prediction error)^4, 5^. Prediction errors, in turn, allow adaptation and updating of predictions to continuously optimize behavior.

In audition these processes have been studied in voice production and perception. Neural activity in the auditory cortex (AC) is suppressed when we speak and hear our voice as compared to when we listen to someone else’s voice^6^. This suppression is the result of comparing predicted and actual sensory feedback to own voice. Electrophysiologically, this phenomenon is captured by the N1 event-related potential (ERP) suppression effect, i.e., the difference in the N1 amplitude for self-generated and externally-generated voices during real-time talking^7–9^ but also when own voices are “self-generated” via a button-press^10^. Changes in the acoustic properties of the self-generated voice, for example, during a cold or vocal strain, can result in a mismatch between the predicted and the actual sensory feedback to the own voice. These mismatches reduce the N1 suppression effect and may lead to the allocation of additional attentional resources to sensory feedback processing and to the attribution of higher prominence to the self-generated voice^11–13^. This likely explains why empirical studies have reported both an increased N1 and N2 response in unexpected sensory feedback processing^14–18^. Unexpected changes in sensory feedback might evoke a surprise response (increased N1^12, 13^) that, in turn, can increase error awareness and attentional control (increased N2^19, 20^).

### Hallucination proneness and sensory suppression

Auditory verbal hallucinations (AVH) can occur in healthy individuals with a prevalence of 6-13%,^21, 22^ implying a continuum of proneness ranging from no to infrequent or frequent AVH experiences^23–27^. Altered sensory feedback processing, resulting from insufficient monitoring or inaccurately attributing the self-generated voice to an external source, likely forms the core of AVH experiences^28–34^. These alterations were reported for voice hearers with a psychotic disorder and non-clinical voice hearers, suggesting that aberrant sensory feedback processing in own voice production is a common feature associated with AVH, regardless of the clinical status of the participant^28, 29, 31, 35–40^. For example, the N1 suppression effect is reduced in playback in participants with increased hallucination proneness (HP^10^), and it is reversed in real-time voice production tasks in persons with a psychotic disorder^7–9, 41, 42^. While the underlying cognitive and neural mechanisms of AVH seem to somehow overlap in voice hearers with and without a psychotic disorder^28, 29, 43, 44^, differences pertain to the perceived emotional quality, appraisal, controllability, and related distress^22, 45^. Unlike non-clinical voice hearers, voice hearers with a psychotic disorder often experience negative, derogatory, and life-threatening voices^21, 46–48^. This distinction in emotional voice quality and the potentially resulting distress are linked to deficits in identification and appraisal of vocal emotions in both voice hearers with ^49–51^ and without a psychotic disorder^52^. Unlike non-clinical voice hearers, voice hearers with a psychotic disorder not only tend to ascribe more attention to negative emotions and perceive them more strongly but also misattribute negative meaning to neutral stimuli to maintain positive symptoms such as AVH^53–56^. Misattributions of salience and the source of a self-generated stimulus in voice hearing were linked to aberrant predictive processing^33, 34, 57–59^. Abnormally strong top-down predictions might generate attentional biases, causing an imbalance between expected and actual sensory input^60–63^. This imbalance might lead to difficulties in perception, for example, the misattribution of negative meaning to neutral stimuli and perceiving meaningful information (e.g., speech) in noise^60, 63–66^, ultimately leading to false perceptions - AVH. Taken together, these findings emphasize the interdependence and mutual influence between alterations in sensory perception and predictive processing in voice hearers. Therefore, by manipulating emotional quality and thereby altering the perceptual certainty of recognizing one’s own voice, one can probe both changes in sensory predictive processing as well as attention allocation in high HP persons, highlighting transitions along the HP spectrum.

### The current study and rationale

Using a well-validated EEG motor-auditory task and building on own prior work (Figure 1)^10^, the current study examined whether systematic changes in sensory feedback processing of the self-voice as a function of HP lead to altered sensory suppression (N1, P2) and attentional control (N2). The emotional quality of the self-voice was manipulated to change the level of certainty in sensory feedback processing (100% neutral, 60-40%: neutral-angry; 50-50%: neutral-angry; 40-60%: neutral-angry and 100% angry). For the self-voice that is most certain (100% neutral and 100% angry), we expected a reduction of the classical N1 suppression effect (self- < externally-generated) with higher HP^10^. For the uncertain self-voice (60-40%: neutral-angry; 50-50%: neutral-angry; 40-60%: neutral-angry), we expected a reversed N1 suppression effect (self- > externally-generated) with increasing levels of uncertainty regarding sensory feedback, in persons with low compared to high HP. Similar effects were expected for the P2 response that indicates the conscious detection of self-generated stimuli^15, 67, 68^. Considering that the presumed alterations are linked to attentional control and error awareness, a reduced or reversed N2 suppression effect (self- > externally-generated) was expected for the certain compared to uncertain self-voice with higher HP.

**Figure 1:**
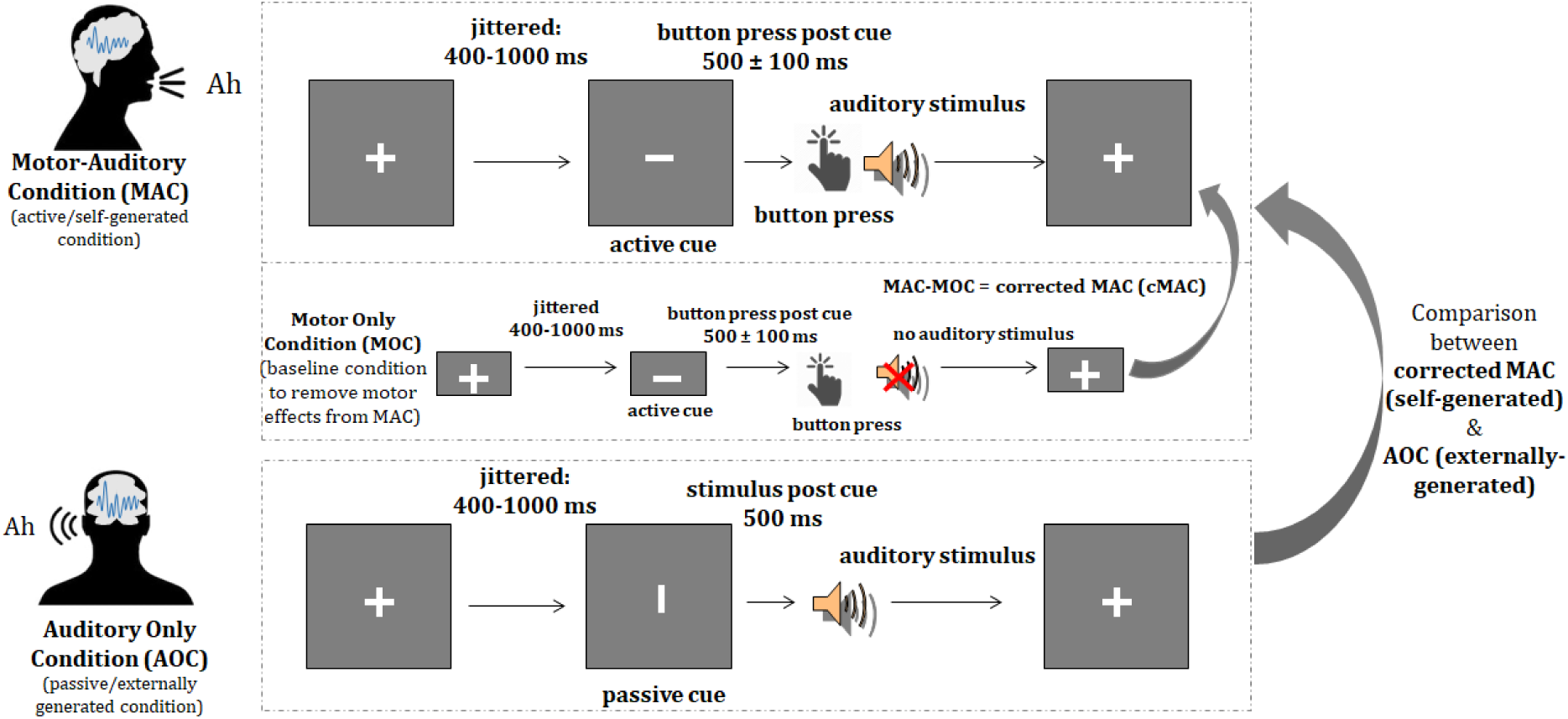
Graphical representation of the Motor-auditory task. Abbreviations: MA = Motor Auditory Condition; AO = Auditory Only Condition; MO = Motor Only Condition. Motor activity from MA condition was removed by subtracting MO from MA to obtain MA corrected condition. Statistical analyses were performed with ERPs from MAc and AO conditions.

## Methods

### Participants

Twenty-nine healthy adults (age range 18-27 years) were recruited. All participants were first invited for a voice recording, followed by the EEG session. Three participants did not participate in the EEG sessions due to time constraints, whereas one participant was excluded from further analysis due to technical issues during the EEG data collection. Therefore, the final participant number was 25 (21 females, mean age = 21.24, s.d. = 2.49 years; 21 right-, 3 left-handed, and 1 ambidextrous) varying in HP (Launay Slade Hallucinations Scale (LSHS)^69–72^ total scores: mean = 18.56, s.d. = 10.17, max = 42, min = 3; LSHS AVH scores [sum of items: “In the past, I have had the experience of hearing a person’s voice and then found no one was there”, “I often hear a voice speaking my thoughts aloud”, and “I have been troubled by voices in my head”]: mean = 2.40, s.d. = 2.62, min = 0, max = 11). All participants provided their written informed consent before the start of the study. They either received financial compensation (vouchers) or study credits for their participation. All participants self-reported normal or corrected-to-normal visual acuity and normal hearing. The study was approved by the Ethics Committee of the Faculty of Psychology and Neuroscience at Maastricht University and performed in accordance with the approved guidelines and the Declaration of Helsinki (ERCPN-176_08_02_2017_S2).

### Procedure

All participants underwent two study sessions conducted on separate visits. During the first voice recording session, “ah” and “oh” vocalizations from each participant were recorded and morphed (see section A of the supplementary document) to create final (100% neutral, 60-40%: neutral-angry; 50-50%: neutral-angry; 40-60%: neutral-angry and 100% angry) voice morphs for the EEG experiment. During the second session, EEG was recorded while the participants performed the auditory-motor task (Figure 1; see section A of the supplementary document). The task was programmed and presented using the Presentation software (version 18.3; Neurobehavioral Systems, Inc.). Stimuli were presented via ear inserts. Button presses were recorded via the spacebar button on the keyboard. Participants were given an overview of the procedure and the principles of EEG at the start of the session. They sat comfortably in an electrically shielded soundproof chamber in front of a screen placed about 100 cm away. Participants filled in the LSHS questionnaire while the EEG cap was prepared.

The paradigm was presented in a fully randomized event-related design over 12 runs. Each run consisted of 80 trials (40 AO, 40 MA, and 10 MO). Each trial started with a fixation cross, after which the presentation (vertical or horizontal) of a cue was jittered between 400-1000 ms. The cue was then followed by an auditory stimulus (after 500 ms for AO) or a button press that may (MA) or may not (MO) elicit an auditory stimulus. Five types of voice morphs consisting of “ah” and “oh” vocalizations, respectively, were presented in the AO and MA conditions. Thus, each run consisted of 4 trials of 10 stimulus types each (“ah” and “oh” for 5 voice morphs). This included 96 trials per voice morph (“ah” and “oh” combined, supplementary table 1). Participants were given short breaks after each run. To minimize potential influences of lateralized motor activity, participants were asked to switch their response hand every three runs. Prior to the experiment, participants were trained to press the button within 500 ± 100 ms after the cue (horizontal bar) to align the presentation of auditory stimuli in the MA and AO conditions and to avoid overlap of cue-elicited and motor activation.

### Stimulus Rating

At the end of the EEG session, participants rated their voices for arousal and valence (supplementary figure 1). They additionally rated the voices on perceived ownness, meaning how much they identified their own voice on a Likert scale (0-10). This was done to ensure that participants recognized their own voice and perceived the emotion expressed by it. Participants were debriefed after the experiment was finished.

### EEG data acquisition and preprocessing

EEG data were recorded with BrainVision Recorder (Brain Products, Munich, Germany) using an ActiChamp 128-channel active electrode set-up while participants performed the auditory-motor task. Data were acquired with a sampling frequency of 1000 Hz, an electrode impedance below 10 kΩ, using TP10 as online reference. During the EEG recording, participants were seated in a comfortable chair about 100 cm away from the screen in an acoustically and electrically shielded chamber.

EEG data were pre-processed (see section A the supplementary document) using the Letswave6 toolbox (https://github.com/NOCIONS/letswave6) running on MATLAB 2019a. The grand averaged waveforms revealed three ERP components, two negative components peaking at approximately 164 ms and 460 ms respectively and one positive component peaking at 286 ms. As the latencies of the ERP responses varied significantly (supplementary table 2), peak amplitudes as an outcome measure were chosen for data quantification. The N1 peak amplitude was defined as the largest negative peak occurring between 80-230 ms, the P2 peak amplitude was defined as the following positive peak between N1 and 380 ms, and the N2 peak amplitude as the negative peak between the P2 and 600 ms^73, 74^. Previous research showed that the ERP components of interest all have prominent fronto-medial and fronto-central topographies^6, 75, 76^. Therefore, the N1, P2, and N2 responses were extracted from the same fronto-central region of interest (ROI) that included 21 electrode locations: AFF1h, AFF2h, F1, Fz, F2, FFC3h, FFC1h, FFC2h, FFC4h, FC3, FC1, FCz, FC2, FC4, FCC3h, FCC1h, FCC2h, FCC4h, C1, Cz, C2 (Figure 2).

**Figure 2:**
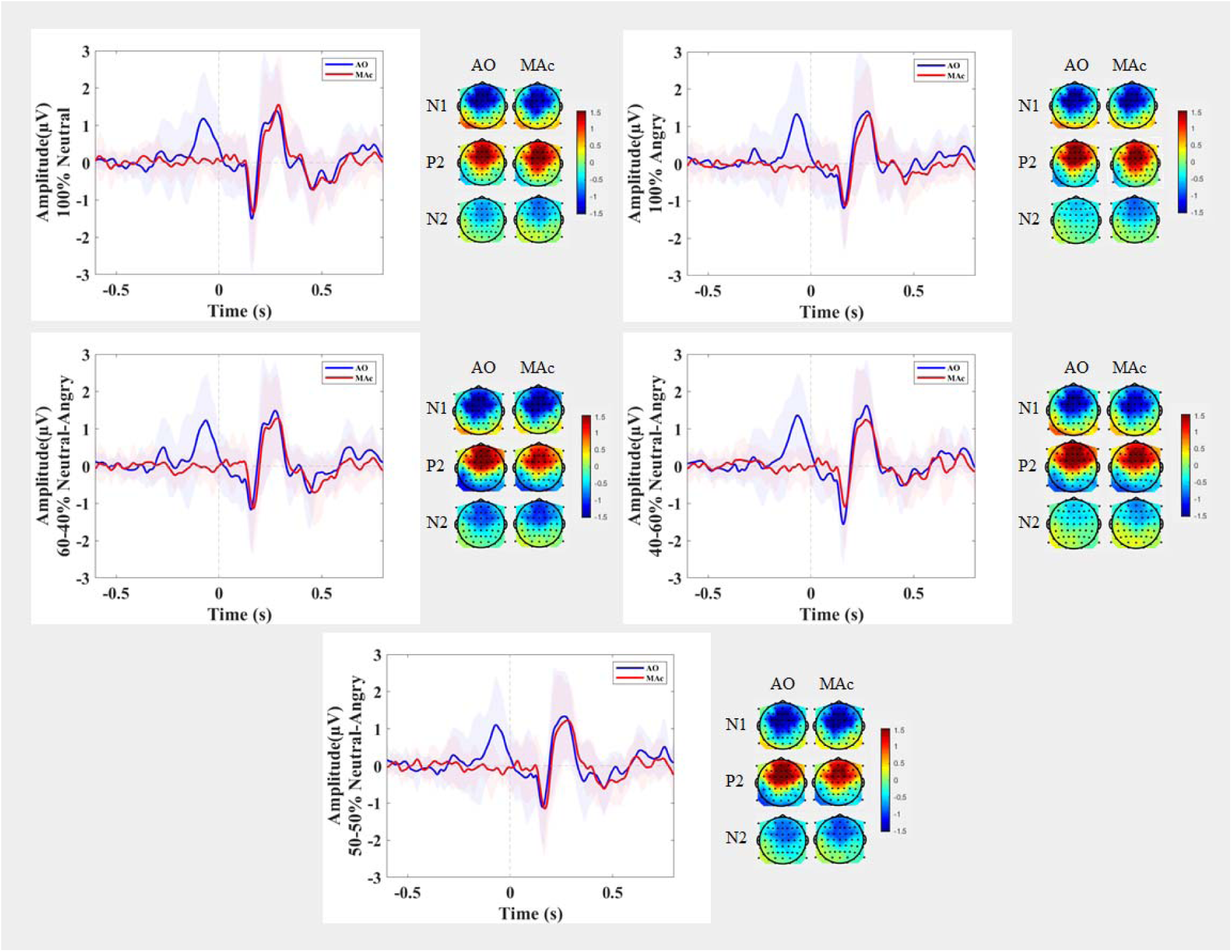
Grand average ERP waveforms ± standard error of mean and topographic maps comparing self-generated and externally-generated voices for the five own voice types originating from a fronto-central ROI. Abbreviations: MAc = Motor Auditory Corrected; AO = Auditory Only.

### Statistical analyses

Statistical analyses on N1, P2, and N2 data were performed in R version 4.2.2 (2022-10-31) Copyright (C) 2022, using linear mixed modeling with lmer and lmerTest packages^77, 78^. We used linear mixed modeling to control for the random effects of participants influencing the outcome measure. Additionally, since HP measured by the LSHS is a continuous variable, Linear mixed modeling was considered more appropriate than classical ANOVA to analyze the impact of HP on sensory feedback (condition) and voice quality (stimulus type). Amplitude values of the ERPs (N1/P2/N2) were used as outcome measures, while participants were used as random effects, and condition (2 levels: MAc and AO), stimulus type (5 levels: 100% neutral, 60-40% neutral-angry, 50-50% neutral-angry, 40-60% neutral-angry, 100% angry) and LSHS total or LSHS AVH scores (continuous variable) were included as fixed effects in the models. For all models, the Gaussian distribution of model residuals and quantile-quantile plots confirmed their respective adequacy.

## Results

We followed a hypothesis-driven approach to probe changes in voice quality (stimulus type) and sensory prediction (condition) as a function of HP.

*N1:* To probe the influence of HP (based on LSHS total scores) on condition and stimulus type, we tested the model [m1_N1 <-lmer(N1 ∼ + Condition * LSHS total + Stimulus Type * LSHS total + (1|ID), data=data, REML = FALSE)] against the null model [m0_N1], which showed the best goodness of fit and yielded a significant difference (χ^2^(11) = 24.072, p = 0.01243*; AIC = 432.93; Table 1, Figure 3). We thus replicated the N1 sensory suppression effect where externally generated (AO) voices lead to a larger (more negative) N1 response than self-generated (MAc) voices. We also observed an overall decrease (less negative) in the N1 response independent of condition (AO or MAc) with increased HP (LSHS total scores) for the angry compared to neutral voice (Table 1, Figure 3).

**Figure 3:**
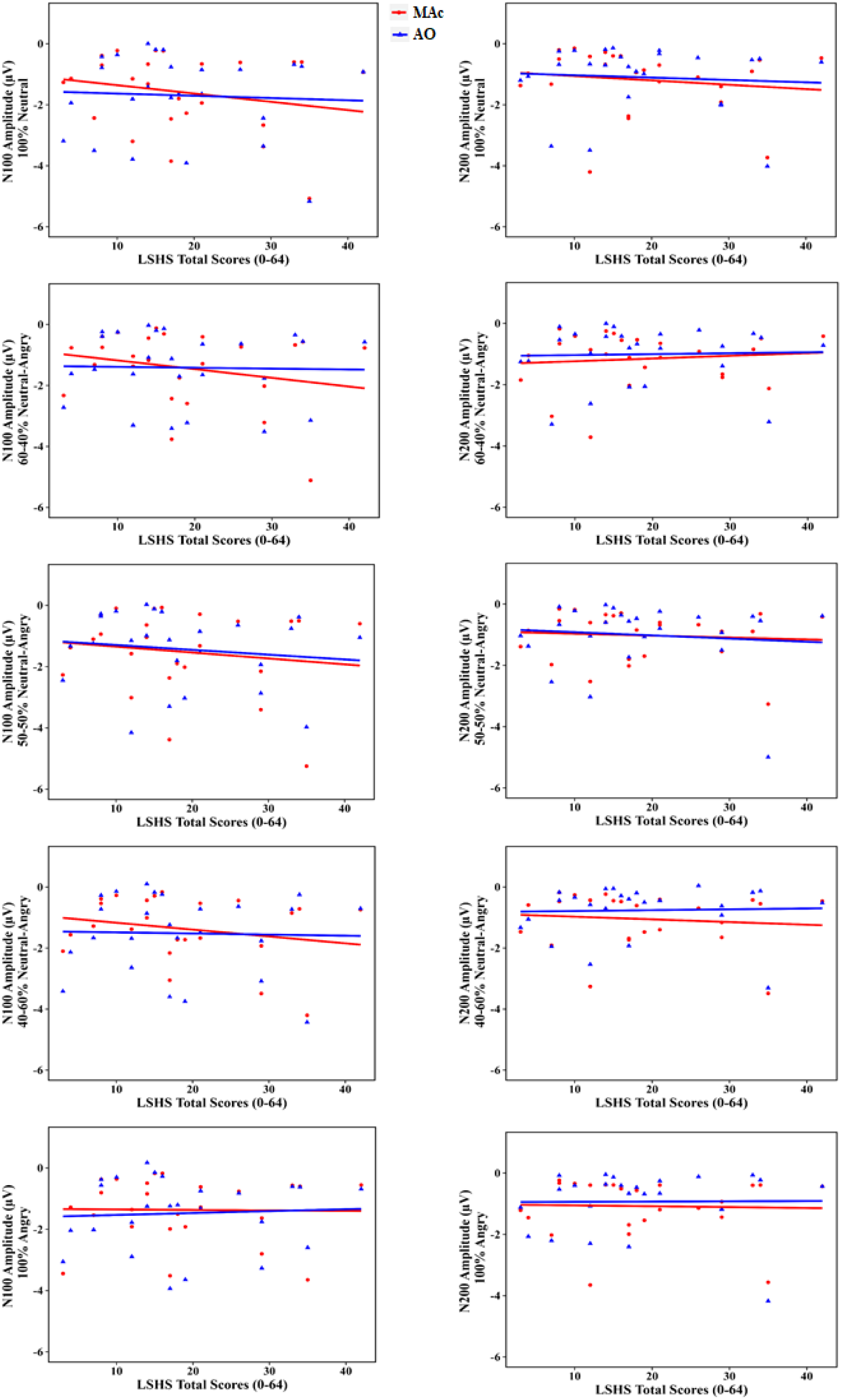
Scatter plots depicting N1 and N2 modulations as a function of HP based on LSHS total scores, for each stimulus type. The N1 response for the self-generated voice increased (more negative) with an increase in HP (Table 1). The N2 response decreased with an increase in HP for the most uncertain self-voice, regardless of the conditions (Table 2).

**Table 1:**
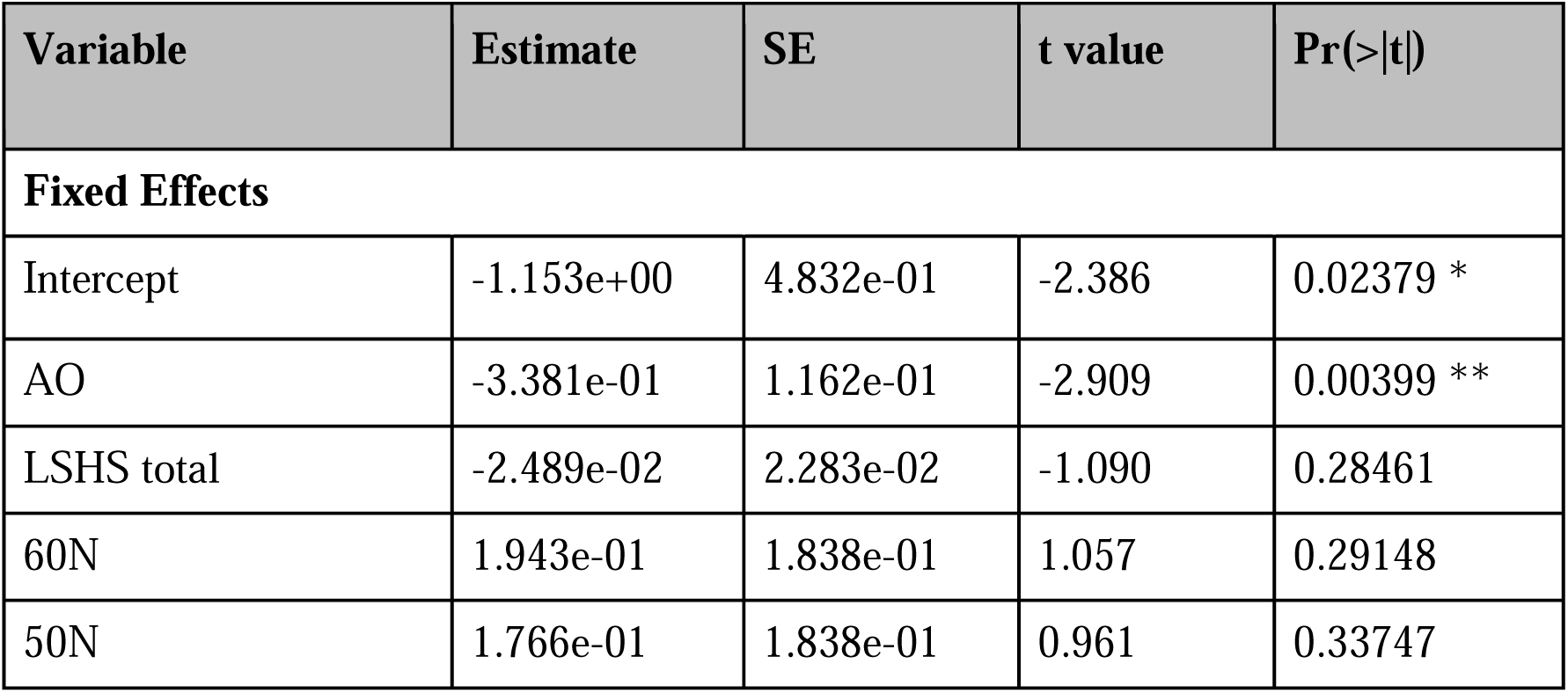
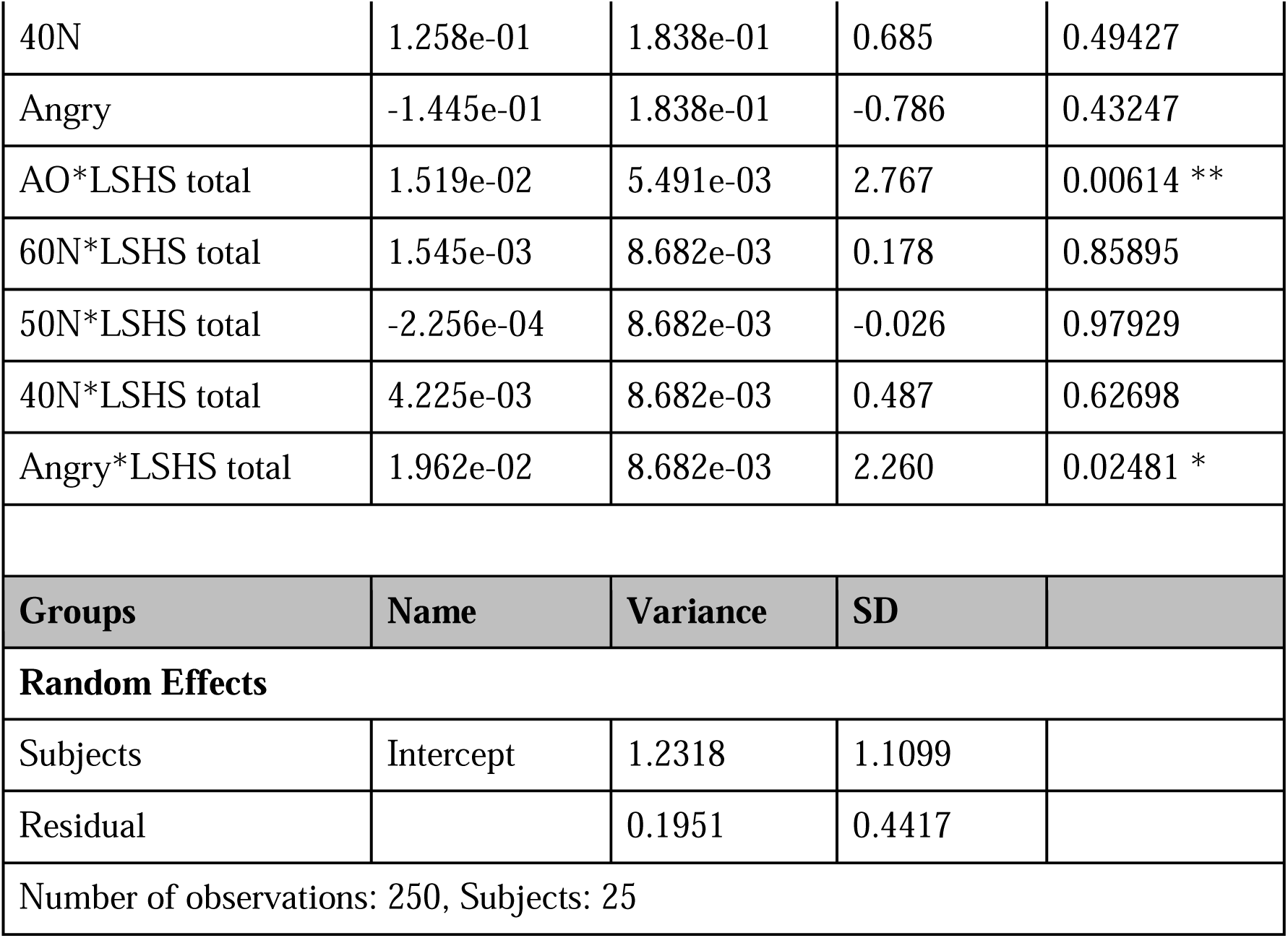
Linear mixed effects model for the N1 including the effect of HP based on LSHS total scores. Abbreviations: SE = standard error; SD = standard deviation; *p < 0.05; **p < 0.01; ***p < 0.001. Degrees of freedom for Fixed Effects: df = 225.0 (except Intercept: df = 29.03).

**Table 2:** Linear mixed effects model for the N2, including the effect of HP based on LSHS total scores. Abbreviations: SE = standard error; SD = standard deviation; *p < 0.05; **p < 0.01; ***p < 0.001. Degrees of freedom for Fixed Effects: df = 225.0 (except Intercept: df = 29.2785).

| Variable | Estimate | SE | t value | Pr(>\|t\|) |
| --- | --- | --- | --- | --- |
| **Fixed Effects** |  |  |  |  |
| Intercept | -0.980059 | 0.378787 | -2.587 | 0.0149 * |
| AO | 0.092415 | 0.093585 | 0.988 | 0.3245 |
| LSHS total | -0.012202 | 0.017896 | -0.682 | 0.5007 |
| 60N | -0.260779 | 0.147970 | -1.762 | 0.0794 . |
| 50N | 0.075303 | 0.147970 | 0.509 | 0.6113 |
| 40N | 0.085665 | 0.147970 | 0.579 | 0.5632 |
| Angry | -0.052273 | 0.147970 | -0.353 | 0.7242 |
| AO*LSHS total | .002340 | 0.004421 | 0.529 | 0.5971 |
| 60N*LSHS total | 0.017018 | 0.006991 | 2.434 | 0.0157 * |
| 50N*LSHS total | 0.002683 | 0.006991 | 0.384 | 0.7015 |
| 40N*LSHS total | 0.008063 | 0.006991 | 1.153 | 0.2500 |
| Angry*LSHS total | 0.010041 | 0.006991 | 1.436 | 0.1523 |

| Groups | Name | Variance | SD |
| --- | --- | --- | --- |
| **Random Effects** |  |  |  |
| Subjects | Intercept | 0.7530 | 0.8678 |
| Residual |  | 0.1265 | 0.3557 |
| Number of observations: 250, Subjects: 25 |  |  |  |

*P2:* Analysis of the P2 followed the same procedure as for the N1. However, the results indicated that HP (based on LSHS total or AVH scores) did not significantly affect sensory prediction (condition) or voice quality (stimulus type) (see section B of the supplementary document).

*N2:* The model that showed the best goodness of fit [m1_N2 <-lmer(N2 ∼ + Condition * LSHS total + Stimulus Type * LSHS total + (1|ID), data=data, REML = FALSE)] also yielded a significant difference (χ^2^(11) = 27.44, p = 0.003941 **; AIC = 323.15; Table 2, Figure 3) when compared against the null model [m0_N2; AIC = 328.59]. The N2 for the self-generated (60N) own voice compared to neutral own voice decreased (less negative) with an increase in HP (LSHS total scores).

## Discussion

This EEG study investigated how changes in sensory feedback processing of the self-voice link to HP and might engage attentional resources by manipulating the emotional quality of the self-voice, thereby altering the certainty of recognizing one’s own voice. The data analyses focused on the N1, P2, and N2 ERP components elicited by the self- and externally-generated [certain (100% neutral, 100% angry) and uncertain (60-40% neutral-angry, 50-50% neutral-angry, 40-60% neutral-angry)] self-voice (Figure 1). The results replicated previous findings^10, 67^, confirming an N1 suppression effect when comparing sensory feedback processing for the self- and externally-generated voice (Table 1). Critically, this N1 suppression effect was reduced in high HP (based on both LSHS total and AVH scores), confirming a link between HP and altered sensory feedback processing (Figure 3). Moreover, regardless of condition, high HP (based on LSHS total scores) was associated with reduced attention allocation indicated by a reduced N1 response to angry compared to neutral voice and lower error awareness demonstrated in a reduced N2 response to the uncertain (60-40% neutral-angry morph) compared to neutral voice (Table 2, Figure 3). However, HP did not modulate the P2 responses. Overall, these results confirm that HP influences sensory feedback processing, and it suggests that attention allocation for the self-generated voice varies with HP in a group of healthy individuals.

### Sensory feedback processing and attention allocation as a function of HP

Replication of the classical N1 sensory suppression effect^10, 14, 15, 67, 68, 79^ (Table 1) likely indicates that the auditory cortex is prepared for the sensory consequences of the self-generated voice. However, increased HP was associated with an increased N1 response for the self-generated voice (Figure 3), thus reducing the N1 suppression effect. This may indicate altered sensory feedback processing for the self-generated voice as well as increased attentional resource allocation towards sensory feedback processing in high HP individuals. One may consider that this alteration and the need for additional resources stem from a less efficient comparison of expected and actual sensory input and the resultant error signal, which might lead to hyper-accentuation of the self-voice. This perspective is supported by previous studies with voice hearers with^8, 9, 42, 80^ and without psychotic disorder ^10^ using similar paradigms. Altered responses to the self-generated voice might indicate that subtle changes in self-monitoring might already be present in healthy persons with high HP.

Furthermore, regardless of condition (AO or MAc), the N1 response to the angry compared to neutral own voice was reduced in high HP participants (Table 1), likely indicating differences in their response when the emotional quality of their voice becomes (fully) negative. Prior research indicates that high HP persons tend to show a dampened negative emotion perception, based on their ability to control attentional bias towards negative cues^53^. Therefore, the current results may point to a link between high HP and reduced appraisal of and inhibition of attention allocation to negative emotions in a non-clinical sample.

Contrary to expectations, HP did not modulate the P2 in sensory feedback processing of the self-voice. The N1 and P2 have been linked to different effects when attributing a sensory event to one’s own action. Whereas the N1 suppression effect seems to reflect the outcome of the comparison of expected and actual sensory input, the P2 was associated with the more conscious realization that a finger tap elicited a related auditory stimulus^14, 15, 68^. The present task, which involved the pseudo-random interweaving of conditions (MA, AO, MO) and stimuli (5 types of “ah” and “oh” vocalizations each), may have precluded sufficient opportunity for the P2 to engage in conscious retrospective processing of a button-press eliciting the self-voice.

The N2 was reduced for the 60-40% neutral-angry compared to the 100% neutral self-voice in high HP individuals regardless of the condition (Table 2, Figure 3). Prior pilot data showed that anger expressed in “ah” vocalizations was already recognized in the initial morphing steps, i.e., the 70-30% neutral-anger voice on the neutral-angry continuum. It is therefore possible that the 60-40% neutral-angry self-voice, among the five presented voice types marks a distinct shift from perceiving something as neutral to detecting anger in the voice imbuing the perception of an uncertain voice. Consequently, this specific self-generated voice may have yielded the most equivocal outcome regarding perceptual uncertainty of the self-voice. Functionally, the N2 has been linked to error awareness, attentional control, and conscious processing of perceptual novelty^81, 82^. Thus, the reduced N2 to this uncertain self-voice in high HP individuals might suggest an altered response to unexpected change or error awareness. Additionally, the N2 has been linked to heightened emotional reactivity to negative rather than neutral stimuli^83^. Taken together, the reduced N2 in high HP individuals may thus indicate down-regulation of negative emotional reactivity, reduced error awareness, and processing of an uncertain self-voice.

Although the N1 suppression effect was observed for the self-generated voice (Table 1), there was no significant interaction between condition (AO, MAc) and stimulus type (five types of self-voice). This suggests that the self-voice manipulations were still within the acceptable range of feasible acoustic changes and therefore, we did not find differential suppression effects for the different types of self-voices (supplementary figure 1). Further, the lack of this interaction in the N1 could be the result of stimulus type probability (2:3 for certain: uncertain). Previous studies showed that higher probability and stimulus repetition result in a stimulus-specific memory trace reflected in early auditory processing as a pronounced N1 suppression^84–86^. Taken together, the unexpected self-voices might not have induced sufficiently different perceptions either because they were presented more frequently, or because they did not differ sufficiently in their acoustic profile. Consequently, there was no difference in the N1 suppression effect among the self-voices.

Some specifics of the task design should be noted. Unlike the classical ERP suppression paradigm, where different conditions are presented in a blocked design^10, 14, 15, 68^, here all conditions and stimuli were presented in a fully event-related design. Due to the mixing of conditions, a cue was introduced to indicate whether the participant was required to press a button to generate a self-voice or to passively listen to the self-voice (Figure 1). While this cue was removed from the MA by subtracting the MO condition for the final analysis, it remained present in the AO condition resulting in a pre-stimulus positive potential (Figure 2). Next to the presence of the cue, the duration between the cue and the auditory stimulus was constant (500 ms). Both factors caused the participants to pay close attention and made them anticipate the onset of the voice in the externally generated condition. However, even though the temporal delay was similar in the self- and externally-generated conditions, we observe a significant N1 suppression effect (AO > MAc, Table 1). This could be attributed to a confluence of factors. Studies have reported that it is not the motor-action per se, but the voluntary intention, involving motor planning, to self-generate an action (e.g,, a voice) that leads to sensory suppression^87, 88^. Further, the increased N1 response in the cued listening condition (AO), excluding motor planning, could be attributed to explicit attention allocation to a self-relevant stimulus (e.g., self-voice)^89–93^. Together, the performance of a motor action in the self-generated condition may take away attention from listening to the generated stimulus, which differs from a cued listening condition^94, 95^. These factors together may influence how attentional resources are directed towards diverse sensory input and to the different N1 responses to the self-versus externally-generated voice.

Taken together, the current results link increased HP to changes in sensory feedback processing and attentional engagement to the self-voice in a healthy participant group. Specifically, these findings suggest that the processing of sensory consequences of one’s own actions are attenuated, however, this attenuation decreases with an increase in HP. Further, high HP is associated with reduced attention allocation to the angry compared to neutral voice, demonstrating their ability to effectively manage negative content^53^. The current findings thus support the continuity perspective regarding changes in sensory feedback processing and attention allocation previously reported in voice hearers^8, 42, 80, 96, 97^. Nevertheless, to strengthen this concept, further investigations involving participants across the psychosis continuum, including healthy persons who do not hear voices, voice-hearers with and without psychotic disorders, are warranted.

## Acknowledgements

The authors would like to thank Joseph Johnson for sharing the paradigm code that was adapted and used in the current experiment. We would also like to thank Alexandra Emmendorfer for sharing her scripts that were modified and used for EEG preprocessing as well as for engaging in valuable discussions on data analysis.

## Data availability

The data that support the findings of this study are available from the corresponding author upon reasonable request.

## Author Contributions

SXD, MS, DL, AP, SK conceptualized and designed the experiment, SXD prepared materials, collected and analyzed the data, and wrote the first draft of the manuscript, SXD, MS, LG, DL, AP, SK refined the manuscript. AP, MS, SK procured funding for the project. All authors have approved the final version of the manuscript.

