## supplementary document for "Hallucination proneness alters sensory feedback processing in self-voice production"

**SECTION A: Methods**

**Stimulus generation**

*Voice recording*

Participants comfortably sat inside an acoustically and electrically shielded chamber with the recording equipment, while the researcher sat outside this chamber. Recordings were made using a Rode NTKb microphone powered by a Rode NTK microphone power supply (http://www.rode.com/microphones/ntk) and processed with the Praat software (<https://www.praat.org>). Participants were instructed to repeatedly vocalize “ah” and “oh” in a neutral (no emotion) and in an angry voice. Vowels were chosen to eliminate semantic content ^1-3^. Participants were asked to vocalize the vowels for 500 ms, and were provided with examples to familiarize them with the target duration of the vocalization. This duration was chosen to properly capture the emotionality while maintaining self-voice recognition. The best voice samples were selected once the participants confirmed that they recognized their recorded voice, that the anger intensity was the highest that they could produce, that they perceived no emotion in the neutral recording, and if the vowels were pronounced clearly. Background noise was eliminated from the recordings using Audacity software (<https://audacityteam.org/>) and a Praat script was applied to normalize the intensity to 70 dB. The duration of the final neutral and angry “ah” and “oh” vocalizations for each participant was 500 ms.

*Morphing*

To create voice samples with varying degrees of emotional content, the pre-recorded neutral and angry self-voices for each individual participant were parametrically morphed to create neutral-to-angry and angry-to-neutral continua. These continua consisted of 11 stimuli with a 10% stepwise increase (neutral-to-angry)/decrease (angry-to-neutral) in emotionality along the continuum (see supplementary table 1). Morphing was performed using the TANDEM-STRAIGHT software ^4-6^ running on MATLAB (R2019a, v9.6.0.1072779, The MathWorks, Inc., Natick, MA). For the final EEG experiments, 100% neutral, 60-40%: neutral-angry; 50-50%: neutral-angry; 40-60%: neutral-angry and 100% angry voice morphs were selected. The intermediate voice morphs were selected based on pilot data that revealed that the maximum uncertainty to differentiate a neutral from an angry self-voice fell in the range of 35-65% morphing. The increase in emotional voice quality (as self-voice stepwise changes from fully neutral to fully angry) and manipulations of uncertainty (most certain: 100% neutral and angry; uncertain: 60-40%: neutral-angry; 50-50%: neutral-angry; 40-60%: neutral-angry self-voice morphs) would probe both changes in sensory feedback to the self-voice and attention allocation resulting from these changes.

*Auditory-motor task*

A variant of an established button-press task was employed to investigate differences between self- and externally-generated auditory stimuli ^7^ (Figure 1). This task comprises three conditions: a motor-auditory condition (MAC), where participants pressed a button to generate their pre-recorded voice; an auditory-only condition (AOC), where participants passively listened to their pre-recorded voice; and a motor-only condition (MOC), where they pressed a button but did not hear their voice. This latter condition was used to control for motor activity resulting from the button-press in the MA condition (MAC-MOC = corrected MAC [cMAC]). Previous studies have consistently shown that there is a reduction in the N1 amplitude in response to self-generated sound via a button-press compared to passively listening to the same sound ^8^^,^ ^9^, indicating that button-presses can be used as a motor-act to self-generate a stimulus (for voices see ^10^).

**EEG data preprocessing**

Data were first cleaned to remove false button presses (e.g., trials with button presses during AO), downsampled to 500 Hz, and then bandpass filtered (1-30 Hz). All channels were re-referenced to the average of the mastoid electrodes. Eye blinks and movements and noisy electrodes were removed using an independent component analysis (ICA) with the runica algorithm in combination with Rajan and Rayner (PICA) as implemented in Letswave6 (<https://github.com/NOCIONS/letswave6>). ICs representing noise were removed for each participant based on the IC time course and topography. The resulting data were segmented with a pre-stimulus time window of -600 to 800 ms, time-locked to the onset of the auditory stimulus. The segmented data were baseline corrected to a window of -600 to -400 ms relative to the onset of the respective auditory stimulus. This remote baseline window was selected due to a cue-related ERP modulation before the onset of the auditory stimulus in AO, which could not be removed using high-pass filtering. After baseline correction, an automatic artifact rejection algorithm was applied with an amplitude criterion of ± 65µV to remove epochs/trials with remaining artifacts. The resulting data were then averaged for each participant and each condition.

**SECTION B: Table and table legends**

**Supplementary table 1:** Neutral-Angry continua with 11 voice morphs.

a)	Neutral-to-angry

| Emotion/Morphs | **1*** | 2 | 3 | 4 | **5*** | **6*** | **7*** | 8 | 9 | 10 | **11*** |
| --- | --- | --- | --- | --- | --- | --- | --- | --- | --- | --- | --- |
| Neutral | **100%** | 90% | 80% | 70% | **60%** | **50%** | **40%** | 30% | 20% | 10% | **0%** |
| Angry | **0%** | 10% | 20% | 30% | **40%** | **50%** | **60%** | 70% | 80% | 90% | **100%** |

b)	Angry-to-neutral

| Emotion/Morphs | **1*** | 2 | 3 | 4 | **5*** | **6*** | **7*** | 8 | 9 | 10 | **11*** |
| --- | --- | --- | --- | --- | --- | --- | --- | --- | --- | --- | --- |
| Angry | **100%** | 90% | 80% | 70% | **60%** | **50%** | **40%** | 30% | 20% | 10% | **0%** |
| Neutral | **0%** | 10% | 20% | 30% | **40%** | **50%** | **60%** | 70% | 80% | 90% | **100%** |

c)	Final Stimuli for Ah and Oh vocalizations.

100% Neutral = Ah (a1 + b11) + Oh (a1 + b11)

60-40% Neutral-Angry = Ah (a5 + b7) + Oh (a5 + b7)

50-50% Neutral-Angry = Ah (a6 + b6) + Oh (a6 + b6)

40-60% Neutral-Angry = Ah (a7 + b5) + Oh (a7 + b5)

100% Angry = Ah (a11 + b1) + Oh (a11 + b1)

Note: a1 refers to the specific voice morph from table a, voice morph 1.

**Supplementary table 2:** Latency range of N1, P2 and N2 amplitudes.

|  | Min (ms) | Max (ms) |
| --- | --- | --- |
| N1 | 0.08 | 0.23 |
| P2 | 0.20 | 0.38 |
| N2 | 0.25 | 0.6 |

The influence of proneness to auditory verbal hallucinations on condition and stimulus type was tested based on LSHS AVH scores. The respective model [m2_N1 <- lmer (N1 ~ + Condition * LSHS AVH + Stimulus Type + (1|ID), data=data, REML = FALSE)] showed the best goodness of fit and yielded a significant difference (χ2(7) = 14.071, p = 0.04993*; AIC = 434.94) compared to the null model [m0_N1] (Supplementary table 3, and figure 3). A more negative N1 response was observed for external compared to self-generated voices. The N1 response decreased (i.e., it was less negative) in response to externally-generated compared to self-generated voices with increased HP (LSHS AVH scores). Further, compared to neutral voices, other voices lead to a decreased (less negative) N1 response in the self-generated condition.

**Supplementary table 3:** Linear mixed effects model for the N1 response including the effect of HP based on LSHS AVH scores. Abbreviations: SE = standard error; SD = standard deviation; *p < 0.05; **p < 0.01; ***p < 0.001. Degrees of freedom for Fixed Effects: df = 225.0 (except Intercept: df = 27.632).

| **Variable** | **Estimate** | **SE** | **t value** | **Pr(>\|t\|)** |
| --- | --- | --- | --- | --- |
| **Fixed Effects** | | | | |
| Intercept | -1.47223 | 0.09030 | -4.722 | 6.11e-05 *** |
| AO | -0.1700 | 0.07741 | -2.196 | 0.0291 * |
| LSHS AVH | -0.05934 | 0.08622 | -0.688 | 0.4974 |
| 60N | 0.22298 | 0.09030 | 2.469 | 0.0143 * |
| 50N | 0.17245 | 0.09030 | 1.910 | 0.0574 |
| 40N | 0.20424 | 0.09030 | 2.262 | 0.0247 * |
| Angry | 0.21959 | 0.09030 | 2.432 | 0.0158 * |
| AO*LSHS AVH | 0.04744 | 0.02177 | 2.179 | 0.0304 * |
| **Groups** | **Name** | **Variance** | **SD** |  |
| **Random Effects** | | | | |
| Subjects | (Intercept) | 1.2377 | 1.1125 |  |
| Residual |  | 0.2038 | 0.4515 |  |
| Number of observations: 250, Subjects: 25 | | | | |

**Supplementary table 4:** Model comparisons with P2 response as output and HP, Condition and Stimulus. Abbreviations: SE = standard error; SD = standard deviation; NP = number of parameters; AIC = Akaike information criterion; BIC = Bayesian information criterion; Chisq = chi square; Df = degree of freedom; M1 = Condition * Stimulus type * LSHS total/AVH; M2 = Condition * LSHS total/AVH + Stimulus type * LSHS total/AVH; M3 = Condition * LSHS total + Stimulus type; *p < 0.05; **p < 0.01; ***p < 0.001.

| **Models with LSHS total** | | | | | | | | |
| --- | --- | --- | --- | --- | --- | --- | --- | --- |
| **Model** | **NP** | **AIC** | **BIC** | **Log Lik** | **Deviance** | **Chisq** | **Df** | **Pr(>Chisq)** |
| Null model | 3 | 382.48 | 393.05 | -188.24 | 376.48 |  |  |  |
| M1 | 22 | 405.55 | 483.03 | -180.78 | 361.55 | 14.931 | 19 | 0.727 |
| M2 | 14 | 391.02 | 440.32 | -181.51 | 363.02 | 13.468 | 11 | 0.2638 |
| M3 | 10 | 385.52 | 420.73 | -182.76 | 365.52 | 10.966 | 7 | 0.1401 |
| **Models with LSHS AVH** | | | | | | | | |
| **Model** | **NP** | **AIC** | **BIC** | **Log Lik** | **Deviance** | **Chisq** | **Df** | **Pr(>Chisq)** |
| M1 | 22 | 403.75 | 481.23 | -179.88 | 359.75 | 16.729 | 19 | 0.6082 |
| M2 | 14 | 388.50 | 437.80 | -180.25 | 360.50 | 15.98 | 11 | 0.1419 |
| M3 | 10 | 384.58 | 419.79 | -182.29 | 364.58 | 11.904 | 7 | 0.1038 |

The influence of HP was tested based on LSHS AVH. The model [m2_N2 <- lmer (N2 ~ + Condition * LSHS AVH + Stimulus Type*LSHS AVH + (1|ID), data=data, REML = FALSE)] also yielded a significant difference (χ2(11) = 22.32, p = 0.022*; AIC = 328.28) from the null model ([m0_N2]; AIC = 328.59) (supplementary table 5, and figure 6).

**Supplementary table 5:** Linear mixed effects model for the N2, including the effect of HP based on LSHS AVH scores. Abbreviations: SE = standard error; SD = standard deviation; **p* < 0.05; ***p* < 0.01; ****p* < 0.001. Degrees of freedom for Fixed Effects: df = 225.0 (except Intercept: df = 29.39).

| **Variable** | **Estimate** | **SE** | **t value** | **Pr(>\|t\|)** |
| --- | --- | --- | --- | --- |
| **Fixed Effects** | | | | |
| Intercept | -1.236426 | 0.246880 | -5.008 | 2.4e-05 *** |
| AO | 0.148456 | 0.061680 | 2.407 | 0.0169 * |
| LSHS AVH | 0.012458 | 0.069441 | 0.179 | 0.8589 |
| 60N | -0.016236 | 0.097525 | -0.166 | 0.8679 |
| 50N | 0.132913 | 0.097525 | 1.363 | 0.1743 |
| 40N | 0.238609 | 0.097525 | 2.447 | 0.0152 * |
| Angry | 0.095991 | 0.097525 | 0.984 | 0.3260 |
| AO*LSHS AVH | -0.005252 | 0.017349 | -0.303 | 0.7624 |
| 60N*LSHS AVH | 0.029712 | 0.027431 | 1.083 | 0.2799 |
| 50N*LSHS AVH | -0.003256 | 0.027431 | -0.119 | 0.9056 |
| 40N*LSHS AVH | -0.001369 | 0.027431 | -0.050 | 0.9602 |
| Angry*LSHS AVH | 0.015870 | 0.027431 | 0.579 | 0.5635 |
| **Groups** | **Name** | **Variance** | **SD** |  |
| **Random Effects** | | | | |
| Subjects | (Intercept) | 0.7517 | 0.8670 |  |
| Residual |  | 0.1294 | 0.3598 |  |
| Number of observations: 250, Subjects: 25 | | | | |

**SECTION C: Figure and figure legends**

**Supplementary figure 1:** Post experiment stimuli rating. A) Arousal rating on a scale of 0-9 for each voice stimulus. B) Valence rating on a scale of 0-9 for each voice stimulus. C) Ownness rating on a scale of 0-10 for each voice stimulus. Vertical bars represent standard error of mean.

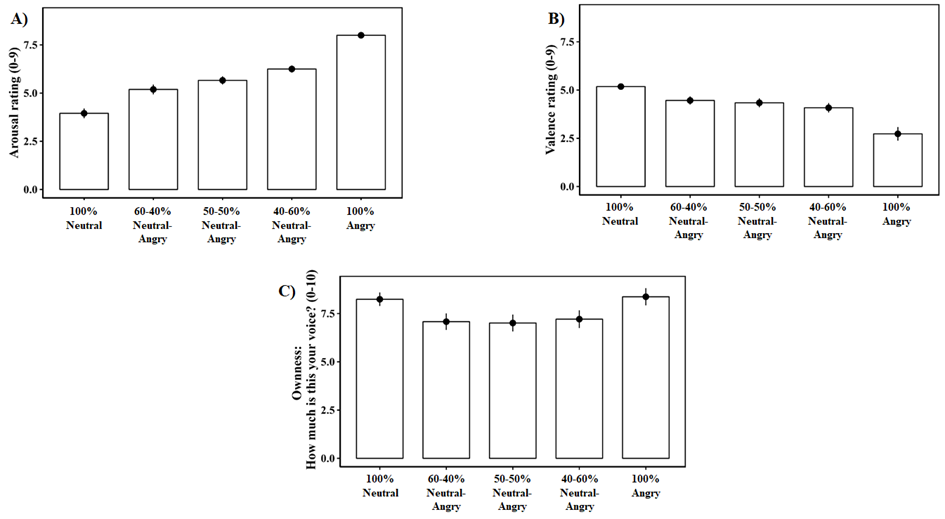

**Supplementary figure 2:** Mean ERP amplitudes for MAc and AO, and suppression effects (AO - MAc) per voice stimulus type. Note: Negative N1 suppression values depict AO > MAc whereas positive N1 suppression values depict MAc > AO. Vertical bars represent standard error of mean.

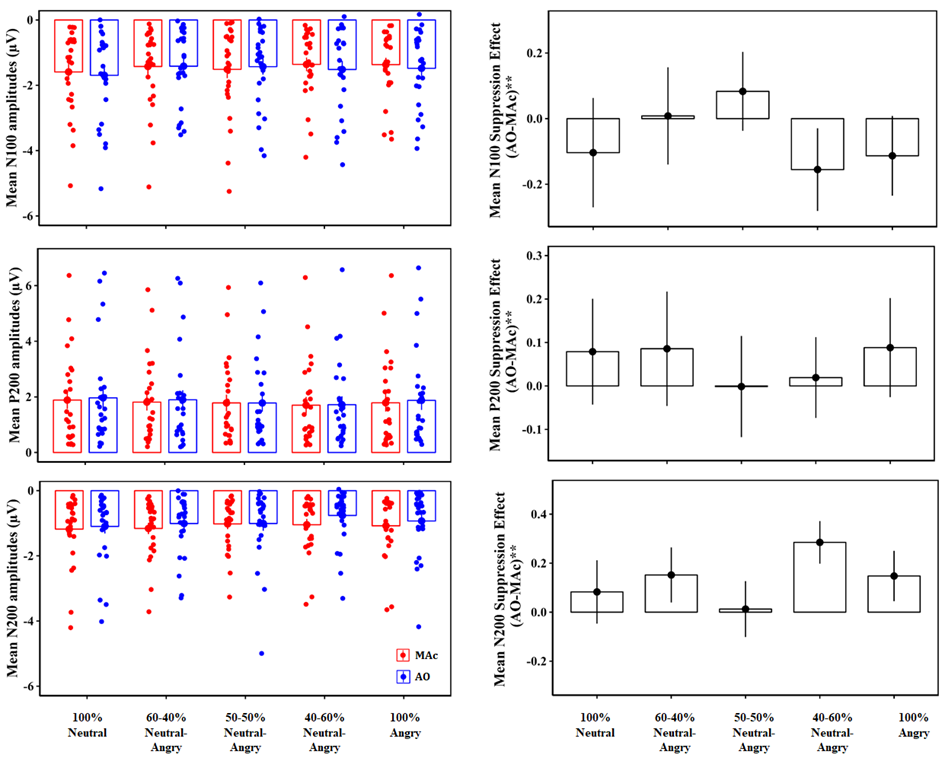

**Supplementary figure 3:** Scatter plots depicting N1 modulation as a function of HP based on LSHS AVH scores for each stimulus type. Increase (more negative) in N1 response for self-generated voice with increase in HP scores (supplementary table 3).

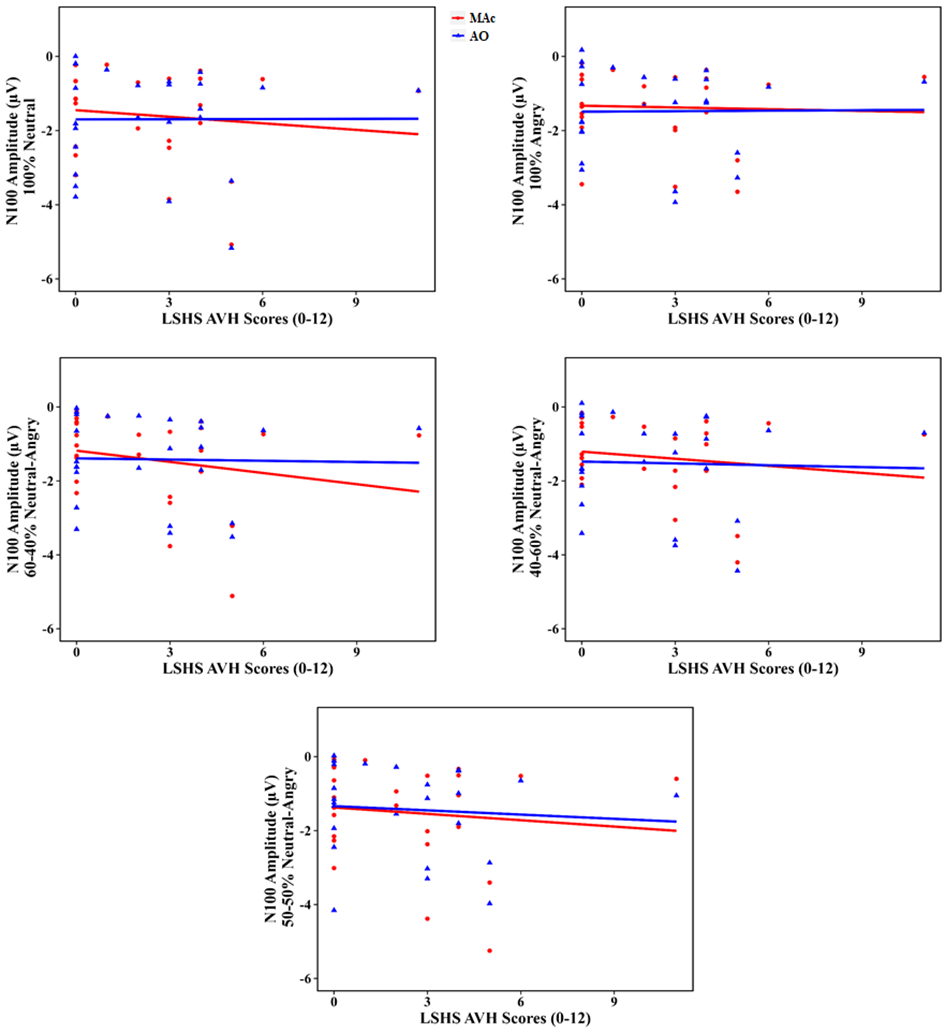

**Supplementary figure 4:** Scatter plots depicting the change in P2 response as a function of HP (based on LSHS total scores) for each stimulus type. No effect of condition (AO, MAc), stimulus type or HP on P2 responses (supplementary table 4).

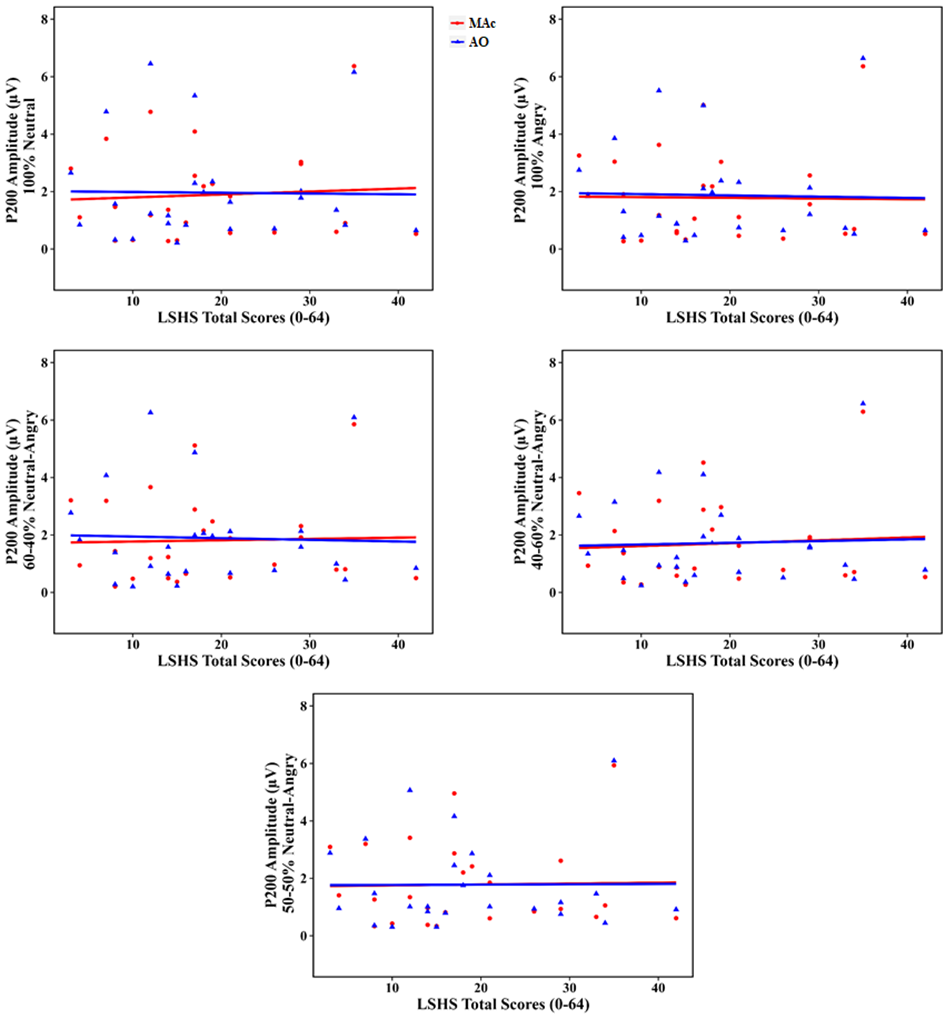

**Supplementary figure 5:** Scatter plots depicting the change in P2 response as a function of HP based on LSHS AVH scores for each stimulus type. No effect of condition (AO, MAc), stimulus type or HP on P2 responses (supplementary table 4).

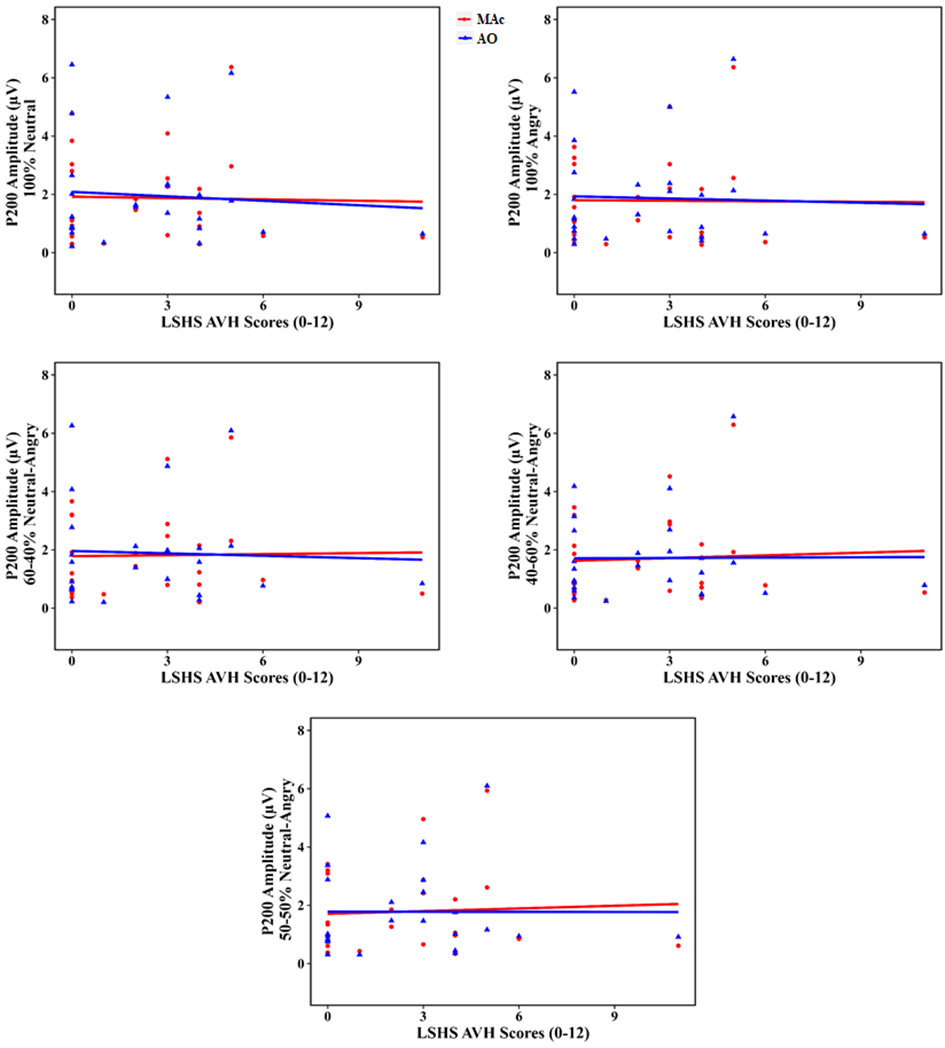

**Supplementary figure 6:** Scatter plots depicting the change in N2 responses as a function of HP based on LSHS AVH scores for each stimulus type (supplementary table 5).

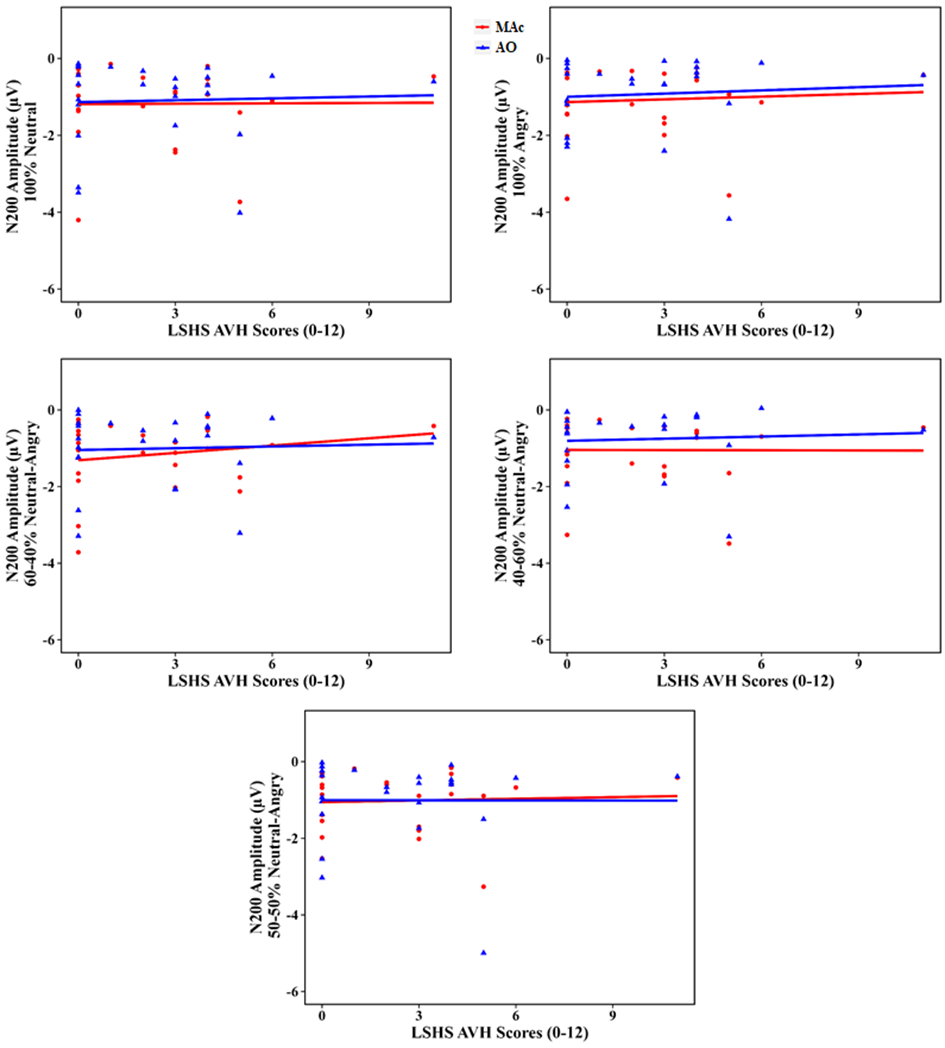
